# Autonomous bioluminescence emission from transgenic mice

**DOI:** 10.1101/2024.06.13.598801

**Authors:** Kamila A. Kiszka, Christian Dullin, Heinz Steffens, Tanja Koenen, Ellen Rothermel, Frauke Alves, Carola Gregor

## Abstract

The bacterial bioluminescence system can be applied to produce autonomous bioluminescence in mammalian cells. Until now, the system has only been stably inserted into cells cultured *in vitro*. Here we report the generation of an autobioluminescent transgenic mouse line constitutively expressing the genes of the bacterial bioluminescence system, enabling substrate-free *in vivo* luminescence imaging of a living mammal.

## Main

Bioluminescent reporter systems, consisting of a luciferase and its substrate luciferin, are frequently applied for *in vivo* imaging of living animals due to the high specificity of the emitted signal and extremely low background. These reporters are particularly suitable for the investigation of processes which require the simultaneous long-term observation of different body regions, such as development, tumor metastasis or the spreading of infections. Improved luciferin-luciferase pairs have been engineered that enable bioluminescent imaging in freely moving animals with single-cell sensitivity^1^. However, such systems suffer from the requirement for an externally supplied luciferin that can produce background luminescence^2^ and exhibit limited biodistribution^3,4^. The luciferin is usually applied to animals by injection that represents an intervention to the organism and makes the overall imaging procedure invasive. Additionally, the luciferin concentration in the body decreases over time^5^, which impedes quantification of the bioluminescence signal during long-term measurements. As an alternative, the luciferin can be continuously produced by the target cells by applying a genetically encoded bioluminescence system which includes enzymes for luciferin biosynthesis and hence enables sustained light emission without any external substrates. An example for such a fully genetically encoded system is the bioluminescence system from bacteria, which has been successfully transferred into mammalian cells and applied for imaging^6–8^. Despite the functionality of this system in *in vitro* models, it has so far not been used to generate transgenic animals to our knowledge and thus no mouse lines or other mammalian model organisms with autonomous bioluminescence are available up to now.

To stably insert the genes of the bacterial bioluminescence system into the mouse genome, we first created an expression cassette containing codon-optimized versions of the involved six genes (Fig. 1a). Besides the bacterial luciferase (*luxAB*), these genes encode the fatty acid reductase complex (*luxCDE*) and a flavin reductase (*frp*) to provide the substrates for the bioluminescence reaction (Fig. 1b). Based on previous work^8^, we chose a gene arrangement where expression of *luxA*+*luxB*+*frp* and *luxD*+*luxE*+*luxC* is driven by separate CMV promoters, with the three genes in each open reading frame separated by viral 2A sequences (Fig. 1a). Transfection of a single plasmid containing this expression cassette into different cell lines resulted in high bioluminescence emission that was similar to the expression of all six genes from individual plasmids and promoters used previously^8^ (Fig. 1c).

**Figure 1.**
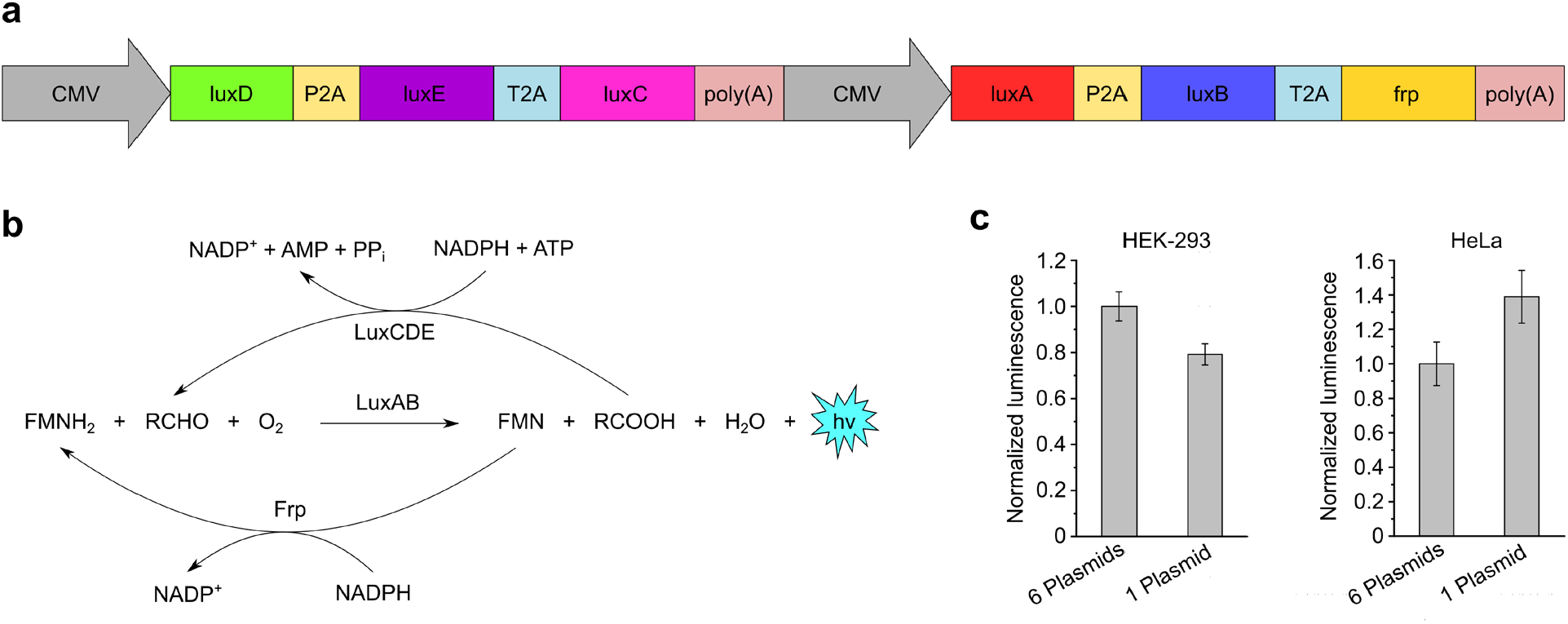
Components of the bacterial bioluminescence system. **a**, Arrangement of the *lux* genes in the expression cassette. **b**, In the bioluminescence reaction, light is generated through the concomitant oxidation of reduced flavin mononucleotide (FMNH2) and a long-chain aliphatic aldehyde (RCHO), catalyzed by the luciferase (LuxAB). The two products are recycled by the fatty acid reductase complex (LuxCDE) and a flavin reductase (Frp). **c**, Comparison of luminescence emission from HEK-293 and HeLa cells transiently transfected with the expression cassette in **a** (1 plasmid) or a mixture of plasmids containing *luxA, luxB, luxC, luxD, luxE* and *frp* separately (6 plasmids). Plots show average values and standard deviations from 5 independent measurements.

We next used the *lux* gene cassette for targeted integration into the *ROSA26* locus of B6(Cg)-Tyr^c-2J^/J mice to generate transgenic animals (hereafter referred to as Lux mice) constitutively expressing all genes of the bacterial bioluminescence system. This albino mouse strain was chosen in order to minimize absorption of the emitted light by the skin and fur. We recorded the bioluminescence of anaesthetized mice with an IVIS Spectrum In Vivo Imaging System (PerkinElmer). Bioluminescence emission was observed throughout the body being strongest at the paws, nose, mouth, eyes and tail (Fig. 2), whereas no luminescence was observed in CDS-negative control mice (Supplementary Fig. 1). The uneven distribution of the bioluminescence emission might on the one hand be caused by different expression levels of the *lux* genes in the respective tissues due to different activities of the CMV promoter^9^. On the other hand, elevated bioluminescence levels may reflect increased metabolic activity since light emission depends on the cellular supply of ATP and NADPH. In addition, part of the emitted light is presumably absorbed by overlying tissue layers and also the fur, making the glabrous regions more visible. The overall distribution of the bioluminescent signal was similar in male and female animals except for a more pronounced luminescence emission from the trunk in male mice (Fig. 2).

**Figure 2.**
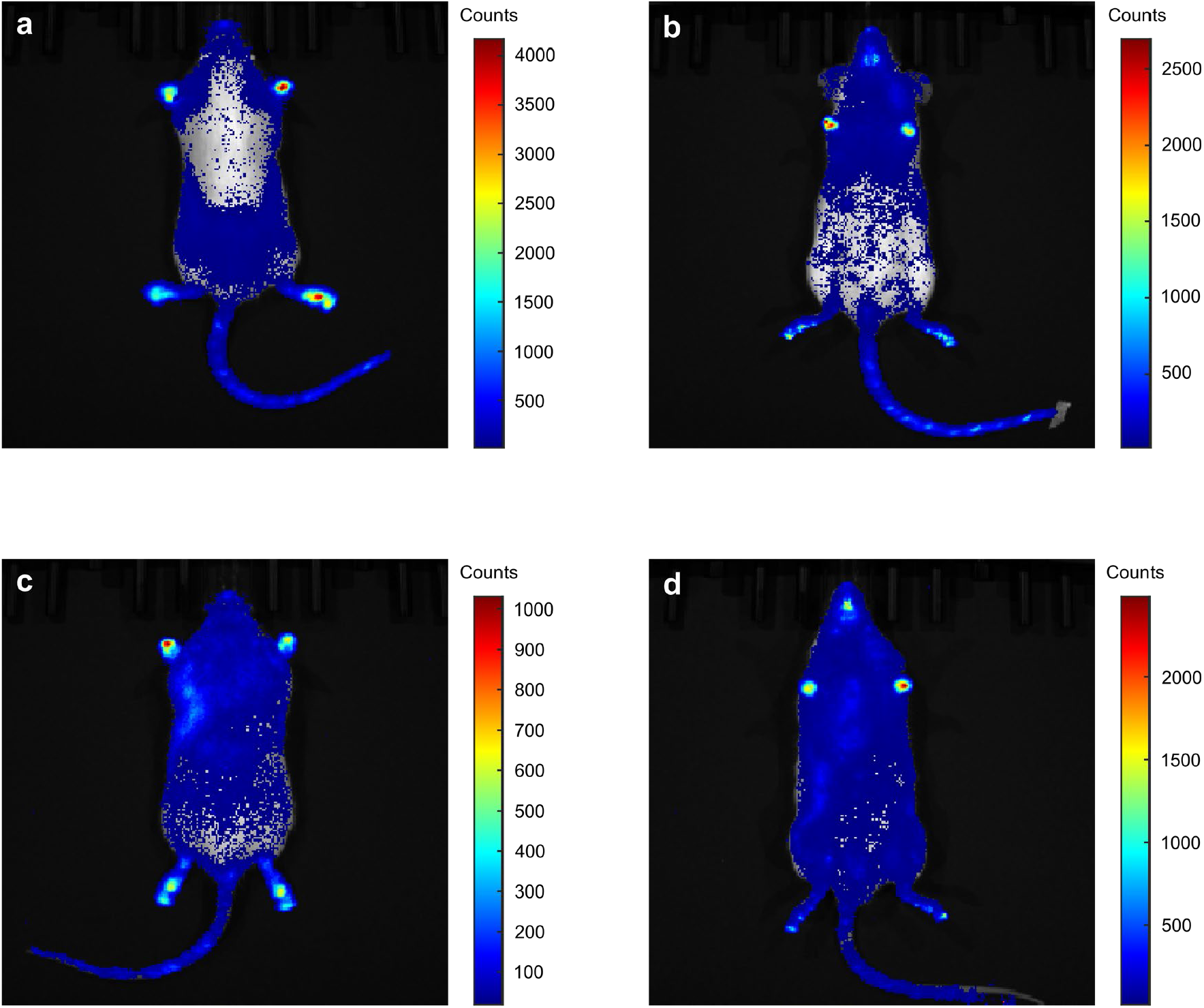
Autonomous bioluminescence emission of living Lux mice. **a, b**, Representative female animal. **c, d**, Representative male animal. Living mice were imaged in prone (**a, c**) and in supine position (**b, d**).

We also imaged bioluminescence of Lux mice after death. Since the generation of bioluminescence requires oxygen, ATP and NADPH, light emission is expected to decrease within seconds to minutes after interruption of the blood supply^10,11^. In line with these expectations, we observed a decrease of the bioluminescence signal in most regions of the animal immediately after death, especially in the tail (Supplementary Figs. 2 and 3). However, light emission persisted for several hours (Supplementary Fig. 3), indicating a long survival time and metabolic activity of the cells despite the lacking blood supply. This may imply that the detected light originates mainly from the skin or directly subjacent tissue, as the upper skin layers are mostly supplied by external oxygen rather than by oxygen transport through the blood^12^. This presumption is supported by the finding that luminescence was present in a piece of skin isolated *post mortem* from the paw of a Lux mouse, along with a reduced luminescence intensity at the corresponding position of sampling (Supplementary Fig. 4). Decrease of the bioluminescence signal over time may therefore be the result of exhaustion of cellular nutrients such as glucose so that cellular energy supply finally becomes limiting for light emission.

By chromosomal incorporation of the genes of the bacterial bioluminescence system, we generated autonomously glowing mice. Since no luciferin supply is required for light emission, such reporter animals are ideal for long-term studies where bioluminescence can for instance be used to monitor gene expression, circadian rhythms and metabolism (e.g., to assess the viability of transplanted tissue or to observe metabolic changes in disease states and upon drug treatment). By further improvements in brightness, we expect that it will be possible in the future to perform such non-invasive measurements also in freely moving animals.

## Methods

### Plasmid construction

Plasmids CMV-luxA-P2A-luxB-T2A-frp-pA pcDNA3.1(+) and CMV-luxD-P2A-luxE-T2A-luxC-pA pcDNA3.1(+) were generated as described in ^8^. The plasmid CMV-luxD-P2A-luxE-T2A-luxC-pA-CMV-luxA-P2A-luxB-T2A-frp-pA pcDNA3.1(+) was generated in the following way: first, the multiple cloning site (MCS) of pcDNA3.1(+) was modified by insertion of the annealed 5’-phosphorylated primers MCS fwd and MCS rev (for primer sequences, see Supplementary Table 1) into pcDNA.1(+) that was previously digested with EcoRI and XhoI.

In the next step, three different inserts were produced. For the first insert, a PCR of luxC was performed using the primers luxC ApaI fwd and luxC AgeI rev. The PCR product was digested with ApaI and AgeI. The second insert was obtained by digestion of the plasmid CMV-luxD-P2A-luxE-T2A-luxC-pA pcDNA3.1(+) with NheI and ApaI and gel purification of the luxD-P2A-luxE band. For the third insert, pA was PCR-amplified from pcDNA3.1(+) with the primers pA AgeI fwd and pA NotI rev and digested with AgeI and NotI. All three inserts were simultaneously ligated into the pcDNA3.1(+) vector containing the new MCS that was previously digested with NheI and NotI.

To insert luxA-P2A-luxB-T2A-frp into the generated plasmid, the CMV promoter was PCR-amplified from pcDNA3.1(+) with the primers CMV NotI fwd and CMV NheI rev. The PCR product was digested with NotI and NheI. luxA-P2A-luxB-T2A-frp was obtained by digestion of the plasmid CMV-luxA-P2A-luxB-T2A-frp-pA pcDNA3.1(+) with NheI and XhoI and gel extraction of the insert. The CMV and luxA-P2A-luxB-T2A-frp insert were simultaneously ligated into the previously generated CMV-luxD-P2A-luxE-T2A-luxC-pA-pA pcDNA3.1(+) plasmid digested with NotI and XhoI.

### Cell culture and imaging

HeLa and HEK-293 cells were obtained from LGC Standards (cat. no. ATCC-CCL-2) and DSMZ (German Collection of Microorganisms and Cell Cultures GmbH, cat. no. ACC 305), respectively.

Cells were grown at 37 °C in 5% CO2 in DMEM with 4.5 g/l glucose (Thermo Fisher Scientific) supplemented with 10% FBS (Merck), 1 mM sodium pyruvate (Thermo Fisher Scientific), 100 units/ml penicillin and 100 μg/ml streptomycin (Merck). For imaging, cells were seeded into 24-well plates and transfected with a total amount of 500 ng DNA (all constructs in pcDNA3.1(+)) using 1 µl jetPRIME transfection reagent (Polyplus-transfection) per well according to the instructions of the manufacturer. For the mixture of 6 plasmids, a luxA:luxB:luxC:luxD:luxE:frp plasmid ratio of 1:1:3:3:3:1 was used as described previously^8^. Bioluminescence imaging was performed 24 hours post-transfection using an Amersham Imager 680 RGB (Cytiva).

### Animals

Animal procedures described here were carried out in accordance with institutional regulations on animals use in research. Experiments performed on living animals were approved and authorized by the Lower Saxony State Office for Consumer Protection and Food Safety (Niedersächsisches Landesamt für Verbraucherschutz und Lebensmittelsicherheit (LAVES); license number: 33.19-42502-04-23-00310). All mice were housed with a 12 hours light/dark cycle and with food and water *ad libitum*.

### Generation of transgenic mice

Transgenic CMV-luxD-P2A-luxE-T2A-luxC-pA-CMV-luxA-P2A-luxB-T2A-frp-pA knock-in mice (Lux mice) were produced by The Jackson Laboratory. The expression cassette was inserted into the *ROSA26* locus of the B6(Cg)-Tyr^c-2J^/J albino mouse line using Bxb1 serine integrase. Lux mice were maintained as heterozygous by crossing with wild-type B6(Cg)-Tyr^c-2J^/J albino mice. Heterozygous Lux mice were fertile and did not exhibit apparent abnormalities.

Animals were genotyped by PCR using DNA from ear biopsies. Four sets of primers were used for genotyping PCRs (see Supplementary Table 2). To verify the DNA sequence of the complete expression cassette, overlapping fragments ∼1 kb in length were generated by PCR amplification from genomic DNA using the primer pairs shown in Supplementary Table 3, purified from agarose gels and sequenced using a standard sequencing service (Microsynth Seqlab GmbH).

### Bioluminescence imaging of mice

Bioluminescence imaging of living and dead mice was performed with an IVIS Spectrum In Vivo Imaging System (PerkinElmer). Ten weeks old, heterozygous Lux mice of both sexes (4 male and 4 female mice) were used for measurements and their non-mutant siblings (CDS-negative mice) were used as negative controls.

For *in vivo* measurements, mice were anaesthetized with 1.0-2.0% isoflurane (Forene, Abbvie) in a 50/50 mix of oxygen and air delivered through a snout-mask at a rate of 0.8 L/min. The eyes were protected from dehydration by application of ointment (Bepanthen, Bayer) and a thermostatic heating plate was used to maintain mouse body temperature at 37 °C. Bioluminescence signal was collected during 3 min exposure times from the whole mouse placed in prone and supine position.

For post-mortem imaging, anaesthetized mice were sacrificed by cervical dislocation. Immediately afterwards, animals were imaged in prone position. All images were acquired at 37 °C using 3 min exposure times.

## Data Availability

Any data generated in this study are available from the corresponding authors upon reasonable request.

## Acknowledgements

The authors thank Prof. S. W. Hell (MPI for Multidisciplinary Sciences) for support during the initial phase of the project and the animal facility of the MPI-NAT and the UMG for excellent advice and support.

This work was funded by the Deutsche Forschungsgemeinschaft (DFG, German Research Foundation) under Germany’s Excellence Strategy - EXC 2067/1-390729940. This work was funded by the Deutsche Forschungsgemeinschaft (DFG, German Research Foundation) – SFB-1286 – 317475864.

## References

1. Iwano, S., Sugiyama, M., Hama, H., Watakabe, A., Hasegawa, N. et al. Single-cell bioluminescence imaging of deep tissue in freely moving animals. Science 359, 935–939 (2018)

2. Zhao, H., Doyle, T. C., Wong, R. J., Cao, Y., Stevenson, D. K. et al. Characterization of coelenterazine analogs for measurements of Renilla luciferase activity in live cells and living animals. Mol. Imaging 3, 43–54 (2004)

3. Lee, K.-H., Byun, S. S., Paik, J.-Y., Lee, S. Y., Song, S. H. et al. Cell uptake and tissue distribution of radioiodine labelled D-luciferin: implications for luciferase based gene imaging. Nucl. Med. Commun. 24, 1003–1009 (2003)

4. Berger, F., Paulmurugan, R., Bhaumik, S. & Gambhir, S. S. Uptake kinetics and biodistribution of 14C-D-luciferin – a radiolabeled substrate for the firefly luciferase catalyzed bioluminescence reaction: impact on bioluminescence based reporter gene imaging. Eur. J. Nucl. Med. Mol. Imaging 35, 2275–2285 (2008)

5. Inoue, Y., Sheng, F., Kiryu, S., Watanabe, M., Ratanakanit, H. et al. Gaussia luciferase for bioluminescence tumor monitoring in comparison with firefly luciferase. Mol. Imaging 10, 377–385 (2011)

6. Close, D. M., Patterson, S. S., Ripp, S., Baek, S. J., Sanseverino, J. et al. Autonomous bioluminescent expression of the bacterial luciferase gene cassette (lux) in a mammalian cell line. PLoS One 5, e12441 (2010)

7. Xu, T., Ripp, S., Sayler, G. S. & Close, D. M. Expression of a humanized viral 2A-mediated lux operon efficiently generates autonomous bioluminescence in human cells. PLoS One 9, e96347 (2014)

8. Gregor, C., Pape, J. K., Gwosch, K. C., Gilat, T., Sahl, S. J. et al. Autonomous bioluminescence imaging of single mammalian cells with the bacterial bioluminescence system. Proc. Natl. Acad. Sci. U. S. A. 116, 26491–26496 (2019)

9. Cheng, L., Ziegelhoffer P. R. & Yang, N. S. In vivo promoter activity and transgene expression in mammalian somatic tissues evaluated by using particle bombardment. Proc. Natl. Acad. Sci. U. S. A. 90, 4455–4459 (1993)

10. Wagner, S. R. 4th & Lanier, W. L. Metabolism of glucose, glycogen, and high-energy phosphates during complete cerebral ischemia. A comparison of normoglycemic, chronically hyperglycemic diabetic, and acutely hyperglycemic nondiabetic rats. Anesthesiology 81, 1516–1526 (1994)

11. Vrselja, Z., Daniele, S. G., Silbereis, J., Talpo, F., Morozov, Y. M. et al. Restoration of brain circulation and cellular functions hours post-mortem. Nature 568, 336–343 (2019)

12. Stücker, M., Struk, A., Altmeyer, P., Herde, M., Baumgärtl, H. et al. The cutaneous uptake of atmospheric oxygen contributes significantly to the oxygen supply of human dermis and epidermis. J Physiol. 538, 985–994 (2002)

