## Supplementary Information for "Autonomous bioluminescence emission from transgenic mice"

**Supplementary Table 1.** Primers used for plasmid construction.

| Primer name | Primer sequence (5'→3') |
|---|---|
| MCS fwd | AATTCTGCAGATATCCAGCACAGTGGCGGCCGCTCAATGC |
| MCS rev | TCGAGCATTGAGCGGCCGCCACTGTGCTGGATATCTGCAG |
| luxC ApaI fwd | CTTGATGGGCCCATGACCAAGAAGATCAGCTTCATC |
| luxC AgeI rev | TGCTTAACCGGTTCAGGGCACGAACACCAG |
| pA AgeI fwd | CTAGTTACCGGTTCTAGAGGGCCCGTTTAAACC |
| pA NotI rev | GTTATCGCGGCCGCCCATAGAGCCCACCGCAT |
| CMV NotI fwd | GATTGAGCGGCCGCTGACATTGATTATTGACTAGTTATTAATAGT |
| CMV NheI rev | TTAAACGCTAGCCAGCTTGGGTCTCCCTA |

**Supplementary Table 2.** Primers used for genotyping.

| Primer Set | Primer sequence (5'→3') |
|---|---|
| 1 | TGTTGCTGTCCTCGTTGTAG |
| | CAGGAGTGCATCGACATCAT |
| 2 | GTAGTGGTTGCCCATGTAGTAG |
| | GCCCATGACCAAGAAGATCA |
| 3 | GTCGCTCTGAGTTGTTATCAGT |
| | GACAAGACCATGCAGGAGTAC |
| 4 | TTGCGGCTCAGGTACTCGG |
| | AATGCCAATGCTCTGTCTAGG |

**Supplementary Table 3.** Primers used for PCR and sequencing.

| Primer Set | Primer sequence (5'→3') |
|---|---|
| 1 | AATGCCAATGCTCTGTCTAGG |
| | TTCTCCTCGGGCAGGGTCTC |
| 2 | TGAATAGCTAGCATGGAGAACGAGAGCAAGTACAAGAC |
| | CAGCTTCTTGCGGATCTTCTCC |
| 3 | GTTAGTGCTAGCATGACCAGCTACGTGGACA |
| | CTGTCGTTCAGGATGGGCAGGT |
| 4 | CGAAGTGCTAGCATGACCAAGAAGATCAGCTTC |
| | TCCATGCTCGAGTCAGGGCACGAACACCAG |
| 5 | CGCCTGGTGACCTACATCAG |
| | CAGCCACACGGTGTCGAAGC |
| 6 | GTTAGTGCTAGCATGAAGTTCGGCAACTTCCTG |
| | TACACCAGGATCTGCTCGAAGT |
| 7 | TAAGTCGCTAGCATGAAGTTCGGCCTGTTC |
| | TGCAGCTGCTCGCTGGCGATG |
| 8 | CGTTAGGCTAGCATGGTGAAGATCCAGCCCATCCCCACCACC |
| | GTCGCTCTGAGTTGTTATCAGT |

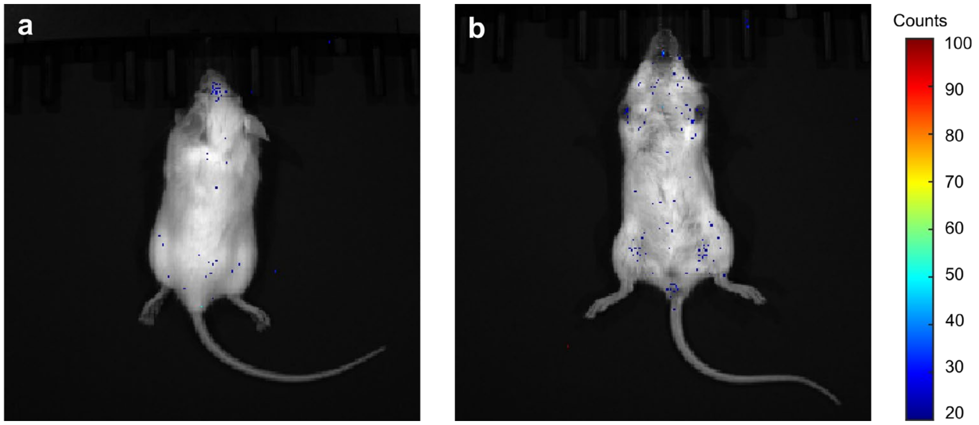


**Supplementary Figure 1. Bioluminescence emission of a living CDS-negative male control mouse.** The living mouse was imaged in prone (**a**) and supine position (**b**).

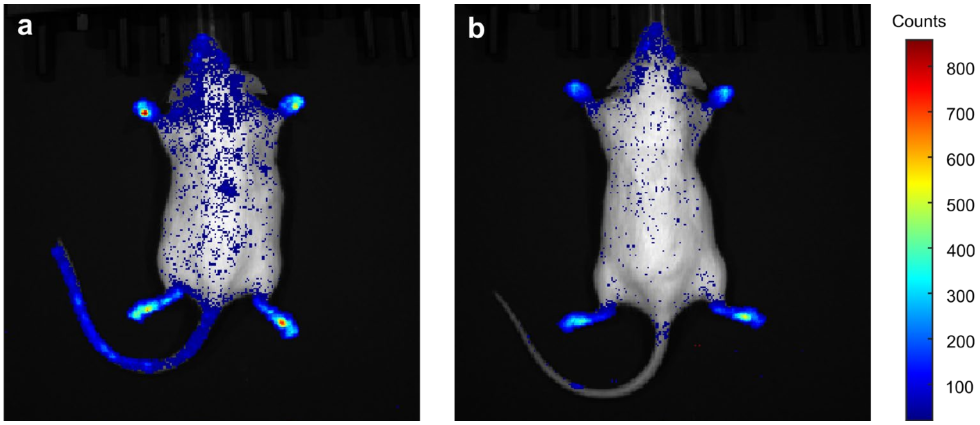


**Supplementary Figure 2. Comparison of bioluminescence emission before and after death.** **a**, Living male Lux mouse. **b**, Same animal as in (**a**), imaged immediately after death.

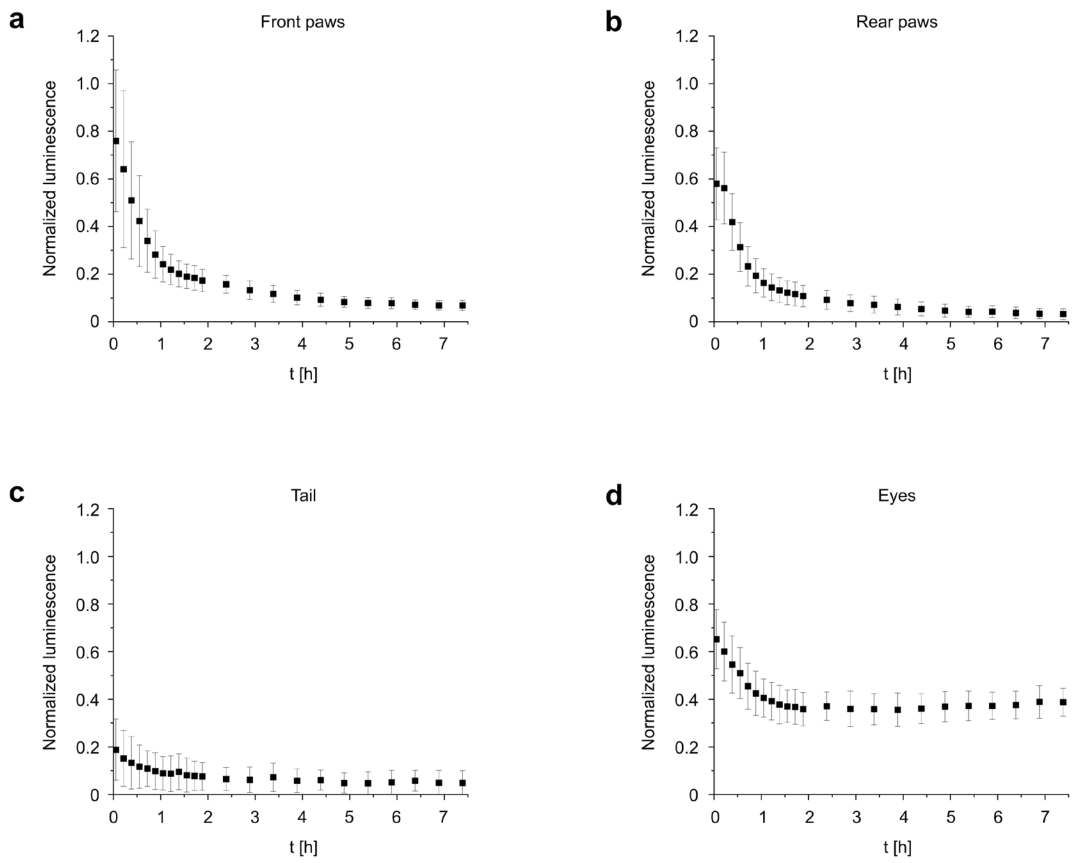


**Supplementary Figure 3. Bioluminescence emission in different regions of Lux mice after death.** The time course of the bioluminescence signal was recorded by the IVIS Spectrum In Vivo Imaging System before and immediately after death in front paws (**a**), rear paws (**b**), tail (**c**) and eyes (**d**). Signals were normalized to the signal of the same region in the living animal. Plots show average values and standard deviations of 5 mice.

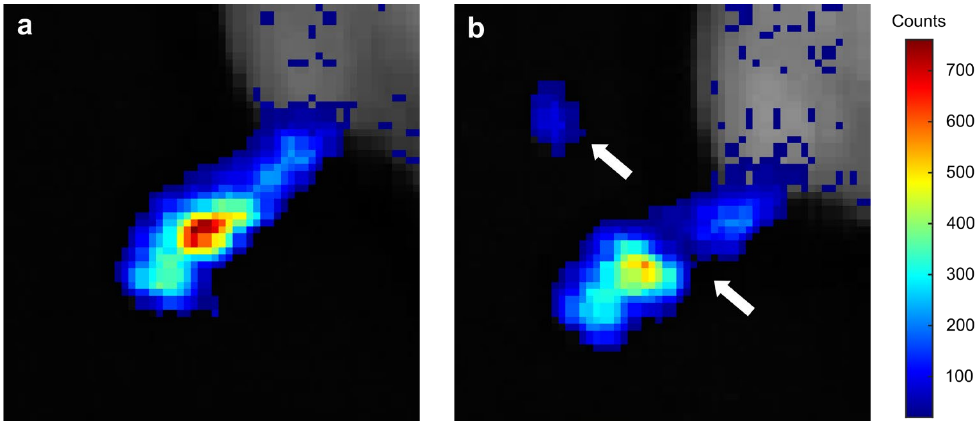


**Supplementary Figure 4. Bioluminescence imaging of isolated skin.** Immediately after death, the mouse was imaged before (**a**) and after (**b**) removal of a piece of skin from a rear paw. The position of sampling and the piece of isolated skin are indicated by arrows.
